# Molecular and functional characterization of the mouse intracardiac nervous system

**DOI:** 10.1101/2021.10.15.464492

**Authors:** Guénaëlle Lizot, Côme Pasqualin, Audrey Tissot, Stephane Pagès, Jean-François Faivre, Aurélien Chatelier

## Abstract

**Background:** The intracardiac nervous system (ICNS) refers to clusters of neurons, located within the heart, which participate to the neuronal regulation of cardiac functions and which are involved in the initiation of cardiac arrhythmias. Therefore, deciphering its role in cardiac physiology and physiopathology is mandatory.

**Objective:** The aim of this study is to provide a phenotypic, electrophysiological and pharmacological characterization of the mouse ICNS, which is still poorly characterized.

**Methods:** Global cardiac innervation and phenotypic diversity were investigated using immunohistochemistry on cleared murine heart and on tissue sections. Patch clamp technique was used for electrophysiological and pharmacological characterization of isolated mouse intracardiac neurons.

**Results:** We have identified the expression of seven distinct neuronal markers within mouse ICNS, thus proving the neurochemical diversity of this network. Of note, it was the first time that the existence of neurons expressing the calcium binding protein calbindin, the neuropeptide Y (NPY) and the cocain and amphetamine regulated transcript (CART) peptide, was described in the mouse. Electrophysiological studies also revealed the existence of four different neuronal populations based on their electrical behavior. Finally, we showed that these neurons can be modulated by several neuromodulators.

**Conclusion:** This study showed that mouse ICNS presents a molecular and functional complexity similar to other species, and is therefore a suitable model to decipher the role of individual neuronal subtypes regarding the modulation of cardiac function and the initiation of cardiac arrhythmias.

## Introduction

Neural control of the heart involves central and peripheral neurons that act interdependently to modulate cardiac parameters such as heart rate, conduction velocity or contractility. As part of this cardiac neuronal regulation, the intrinsic cardiac nervous system (ICNS) is receiving growing attention. Indeed, whereas intracardiac neurons were initially considered as simple parasympathetic postganglionic neurons, studies conducted over the past 30 years have suggested a more complex organization, including sensory, local regulatory and motor neurons within intracardiac ganglia, finally leading to the concept of “little brain on the heart”^1^.

Phenotypic studies have been conducted in many species to identify several neuronal subpopulations. Beside cholinergic phenotype, the presence of catecholaminergic, glutamatergic and nitregic phenotypes have been described among intracardiac neurons^2–10^. This neurochemical diversity has been further confirmed by the expression of several neuropeptides, such as neuropeptide Y (NPY), vasoactive intestinal peptide (VIP), pituitary adenylate cyclase-activating polypeptide, somatotastin (SST), or cocaine and amphetamine regulated transcript (CART) peptide^3,6,10–13^.

From a functional point of view, several studies have also identified various types of cardiac neurons based on their membrane electrical properties^14–17^. In the guinea pig, electrophysiological experiments combined with a morphological characterization of intracardiac neurons led to the identification of three distinct types of neurons, including a putative sensory one^16^. However, the evidence that this neuronal diversity is associated with a functional specialization of neurons is still lacking. Such a characterization would bring essential information to assess the influence of cardiac neuronal network on cardiac physiology. This would be even more important since ICNS have also been implicated in arrhythmias. For example, atrial fibrillation (AF) has been correlated with excessive activity of intracardiac neurons^18^, and specific stimulation of cardiac ganglia has been able to trigger AF^19^. Moreover, it has been showed that ablation of ganglionated plexus reduces AF and is now one of the strategies used in therapy^20^. Deciphering modulation of cardiac function by the ICNS through the role of individual neuronal subtypes is therefore essential but has been for a long time hindered by technical limitations.

Recent advances in genetic engineering have opened new opportunities to improve our understanding of the ICNS. For example, cre-lox systems combined with fluorescent reporter, optogenetic or DREADD approaches represent powerful tools to address precisely the function of one particular subtype of intracardiac neurons. However, because these technologies are almost exclusively available in mouse models, a better understanding of the mouse ICNS is required.

To date, the few existing studies focused on mouse model identified choline acetyltransferase (ChAT), tyrosine hydroxylase (TH) and neuronal nitric oxyde synthase (nNOS) within cardiac neurons^2,7^. However, the expression of other neuronal markers has not yet been investigated.

Moreover, very little is known regarding the electrical and pharmacological properties of mouse cardiac neurons. To our knowledge, only one study investigated this aspect^21^.

This study was therefore designed to better characterize the phenotypic, electrophysiological and pharmacological properties of the mouse ICNS. Immunohistochemical experiments were conducted on cleared whole murine hearts and on tissue sections, allowing us to (1) quantify global autonomic cardiac innervation and (2) investigate the phenotypic heterogeneity of mouse intracardiac neurons. This examination was further strengthened by the characterization of passive and active electrical membrane properties, as well as pharmacological responses of isolated mouse cardiac neurons using the patch clamp technique.

## Methods

### Animals

Experimental procedures were performed using adult C57/BL6 mice (8-20 weeks) of either gender in accordance with the European Union Directive (2010/63/EU) on the protection of animals used for scientific purposes. The protocol was approved by the local ethics committee “COMETHEA”.

### iDISCO heart clearing

Mouse hearts were stained and cleared using a modified iDISCO+ protocol^22^. Images were acquired with the ALICE’s custom-built mesoSPIM microscope at Wyss Center, Geneva. (See supplement).

### Immunohistochemistry

Immunohistochemistry was performed on 40–50 µm heart sections and acquired using a confocal laser scanning microscope (FV3000 Olympus) (See supplement).

### Neuron dissociation

Ganglia were enzymatically digested with 2mL HBSS containing 3mg/mL collagenase type II (Worthington), 7.5mg/mL dispase II and 0.25 mg/mL DNase I (30mn at 37°C). This was followed by an incubation in 2mL trypsin-EDTA 0.25% supplemented with 0.25 mg/mL DNase I (35mn at 37°C). Finally, cells were gently triturated with fire-polished Pasteur pipettes coated with SVF and plated on laminin-coated 35mm Petri dishes.

### Electrophysiology

Electrical membrane properties of isolated cardiac neurons were determined using the whole-cell configuration of the patch clamp technique. Recordings were carried out at room temperature within 30 hours following isolation. Data acquisition and analysis were performed using pClamp software (v11, Molecular devices, San Jose, California, USA).

### Statistical analysis

Statistical analysis was performed using GraphPad Prism (San Diego, California, USA). Data are presented as mean±SEM. Mann-Whitney test was used for statistical comparison. Statistical significance was accepted when p < 0.05.

## Results

### Cholinergic and catecholaminergic innervation of mouse heart

Neural control of the heart involves a combination of peripheral and intrinsic neural structures^1^. In order to investigate global cardiac innervation in mouse, we performed immunodetection of cholinergic (ChAT-IR) and catecholaminergic (TH-IR) structures on cleared murine hearts.

This approach allowed us to appreciate the network of TH-IR nerves innervating mouse hearts (supplementary movie). Large bundles of nerves accessed the heart through the heart hilum, at the base of the heart, and extended toward the dorsal and ventral sides of both ventricles (fig.1a). Ventricular innervation was mainly located at the epicardial surface even if thinner fibers could be seen deeper through the heart wall, especially coursing along the interventricular septum (supplementary movie).

**Figure 1.**
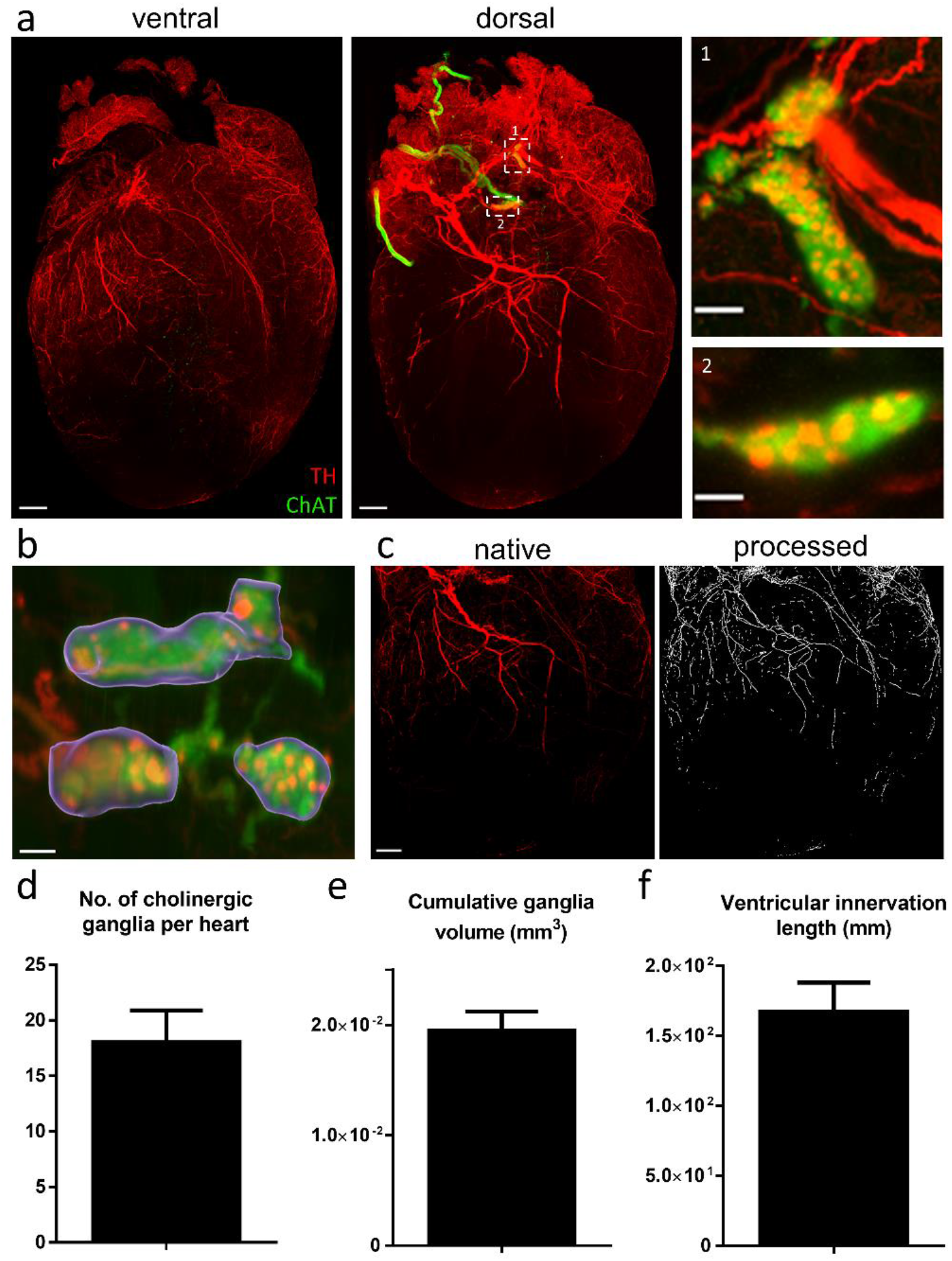
Cholinergic and catecholaminergic cardiac innervation. **a)** 3D projections of the ventral (2500µm z-stack) and dorsal side of a cleared heart (2440µm z-stack) with TH (red) and ChAT (green) staining. Magnification of ganglia are presented in inset 1 and 2. (**b-c)** 3D image processing used for the quantification of ganglia volume (purple) and ventricular innervation (white). **(d-f)** Assessment of the total number of ganglia per heart **(d)**, the cumulative ganglion volume **(e)** and the total length of adrenergic fibers innervating both ventricles **(f)**. Scale bars are 500µm (**a**(ventral and dorsal) and **c**) and 100µm (**a**(insets 1 and 2) and **b**).

ChAT-IR was used to identify intrinsic cardiac ganglia. As seen in figure 1, ganglia were exclusively located in the dorsal side of the heart, in close proximity to the pulmonary veins. The mean number of ganglia per heart was 18±3 ranging from 13 to 23 (n=3, fig.1d). Most of them were characterized by TH-IR puncta, allowing us to see individual neurons (fig.1a insets 1 and 2). Unlike TH, ChAT-IR fibers were almost exclusively located around ganglionated plexus, where they interconnected multiple ganglia.

Global cardiac innervation was further studied by developing a volumetric quantification approach using Imaris. Based on ChAT staining, the volume of each individual ganglion was estimated, resulting in a total ganglion volume of 1.95±0.17 10^−2^ mm^3^ (n=3; fig.1b,e) per heart. The total length of TH-IR nerve fibers innervating both ventricles was quantified using ImageJ software. The total length found was 1.68±0.21 10^2^ mm (n=3; fig.1c,f).

### Neurochemical phenotype of mouse intracardiac neurons

Immunodetection of the cholinergic marker ChAT confirmed the prominent cholinergic phenotype of intrinsic cardiac neurons (98.7%). This marker also labeled intra and interganglionic nerve fibers, as well as varicose terminals surrounding neurons. Immunoreactivity for TH was detected in approximately 30% of cardiac neurons and in numerous nerve fibers (fig.2a-c).

**Figure 2.**
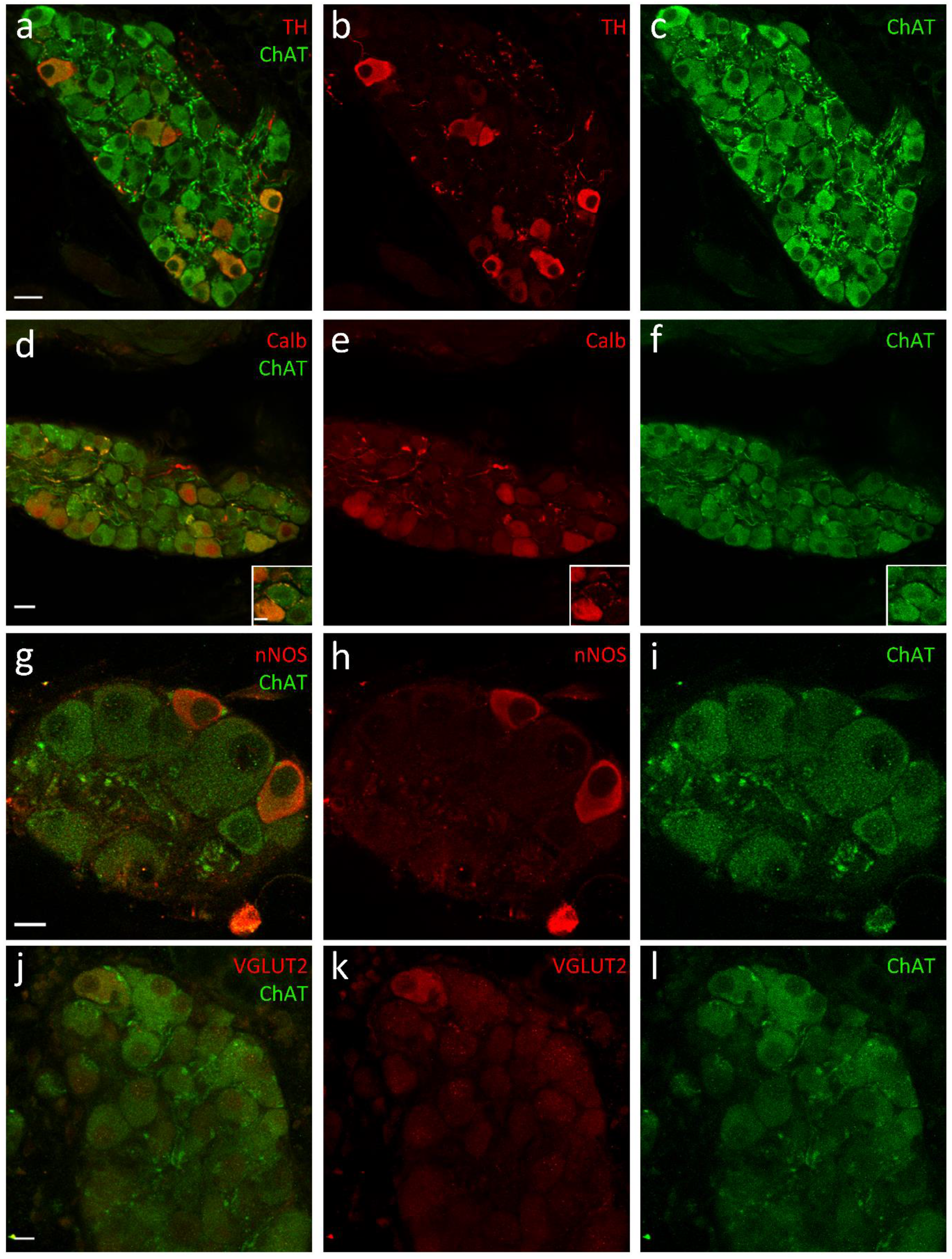
Phenotypic properties of intracardiac neurons. Confocal images of sections of cardiac ganglia immunostained with TH **(a-c)**, calbindin **(d-f)**, nNOS **(g-i)**, VGLUT2 **(j-l)**, and ChAT. Insets in d-f show typical calbindin-IR pericellular baskets surrounding a neuronal somata. Scale bars are 10µm (inset d, g-i and j-l) and 20µm (all others).

Apart from the widely established cholinergic and catecholaminergic markers, we observed that a large proportion of neurons (46%) expressed the calcium binding protein calbindin D-28k (calbindin). In neuronal somata, calbindin-IR was always observed in cytoplasm and often accompanied by a strong nucleus staining (fig.2d-f). Calbindin-IR was also present in multiple intra and inter-ganglionic nerve fibers, and occasionally in pericellular baskets surrounding neuronal somata (fig.2d inset). Calbindin neurons were always co stained by ChAT and 11.8% of cardiac neurons showed immunoreactivity for calbindin, ChAT and TH (Table1).

**Table 1.** Neurochemical profile of mouse intracardiac neurons.

| Neurochemical phenotype | Percentage (number of profiles) | Multiple phenotype (% of total) |
| --- | --- | --- |
| ChAT | 98,7% |  |
| TH | 28,8% (251/873) | 27,9% are ChAT/TH |
| NPY | 66,5% (581/874) |  |
| nNOS | 1,7% (10/577) |  |
| Calb | 45,7% (596/1305) | 11,8% are Calb/ChAT/TH |
| CART | 60,8% (578/950) |  |

A small population of neurons stained with nNOS, the neuronal enzyme responsible for the synthesis of NO, were also present in cardiac ganglia. These neurons were not present in all ganglia and accounted for only 2% of total neurons (fig.2g-i). Even though most nNOS-IR somata coexpressed ChAT, not all of them did.

On very rare occasions, we observed immunoreactivity for the vesicular glutamate transporter 2 (VGLUT2) (fig.2j-l). However, no somata was stained by the other vesicular glutamate transporter VGLUT1. Moreover, despite the use of different markers (Gad67, GABA, TPH2), we did not observe any GABAergic nor serotoninergic phenotype.

Intracardiac neurons have been shown to express several neuropeptides in various species, especially in rodents^6,11^. In mouse hearts, a large proportion of neuronal somata was immunoreactive for NPY (67%) and CART peptides (61%). However, antibody cross-reactions prevented us from quantifying the number of neurons co-expressing both peptides. In somata, both staining were granular with a perinuclear localization, suggesting a localization in the endoplasmic reticulum and the Golgi apparatus (fig.3). Numerous intra and interganglionic nerve fibers were also labeled by these two peptides.

**Figure 3.**
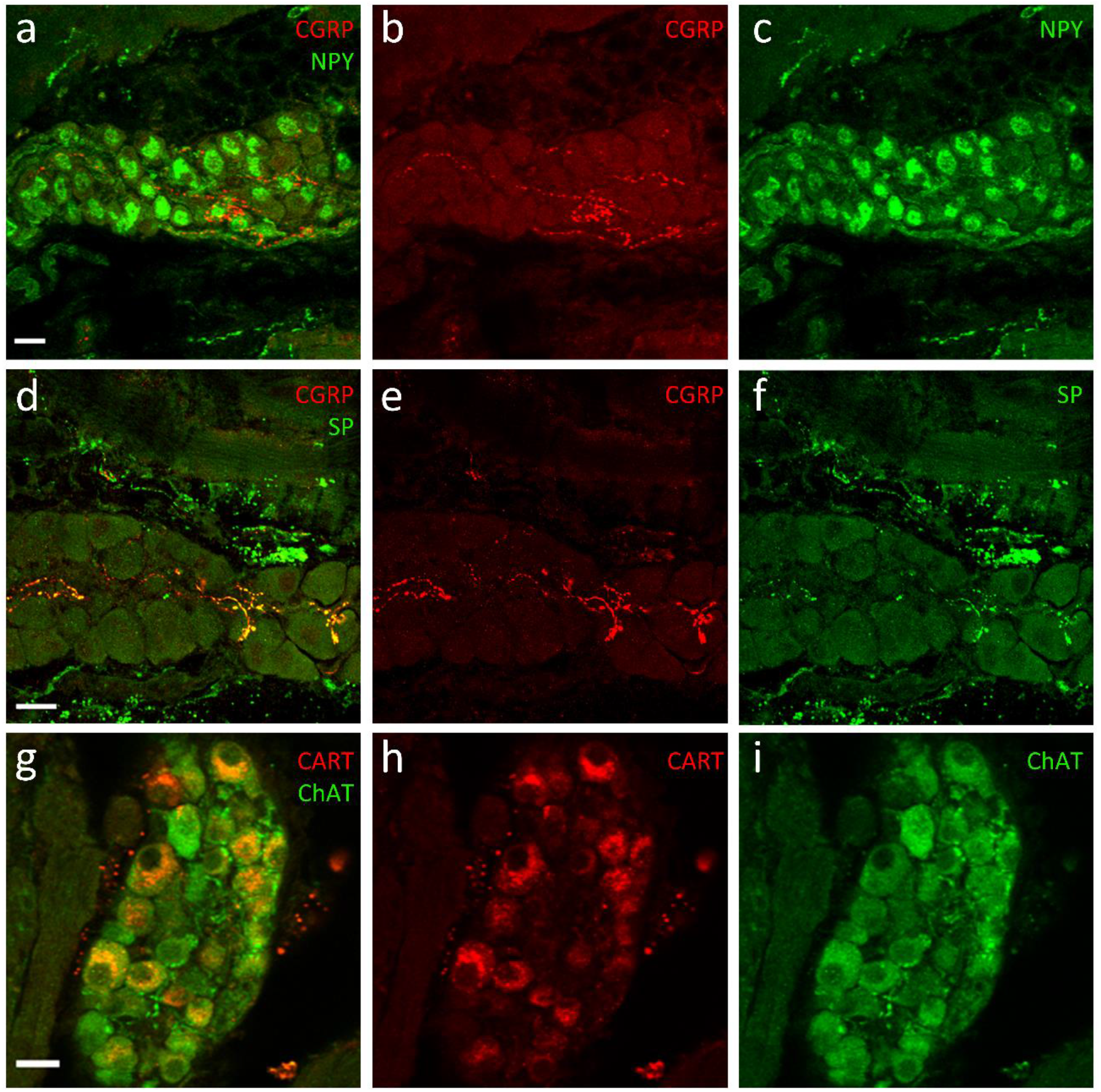
Peptide expression in mouse intracardiac neurons. Confocal images of sections of cardiac ganglia immunostained with NPY and CGRP **(a-c)**, SP and CGRP (d-f) and CART **(d-f)**. Scale bars are 20µm.

Calcitonin Gene Related Peptide-IR (CGRP) and Substance P-IR (SP), two peptides distinctive of sensory neurons, were also observed in nerve fibers, but never in neuronal somata (fig.3a-f). Both peptides were frequently colocalized, although SP-IR nerve fibers were far less numerous. IR for SST and VIP was also tested. However, we did not observe any labeling for these neuropeptides.

### Electrophysiological properties of mouse intracardiac neurons

After isolation, cardiac neurons were identified as cells with rounded cell body with numerous extensions emerging from it (fig.4a). Most neurons were multipolar, even if some uni- and bipolar neurons were also observed.

**Figure 4.**
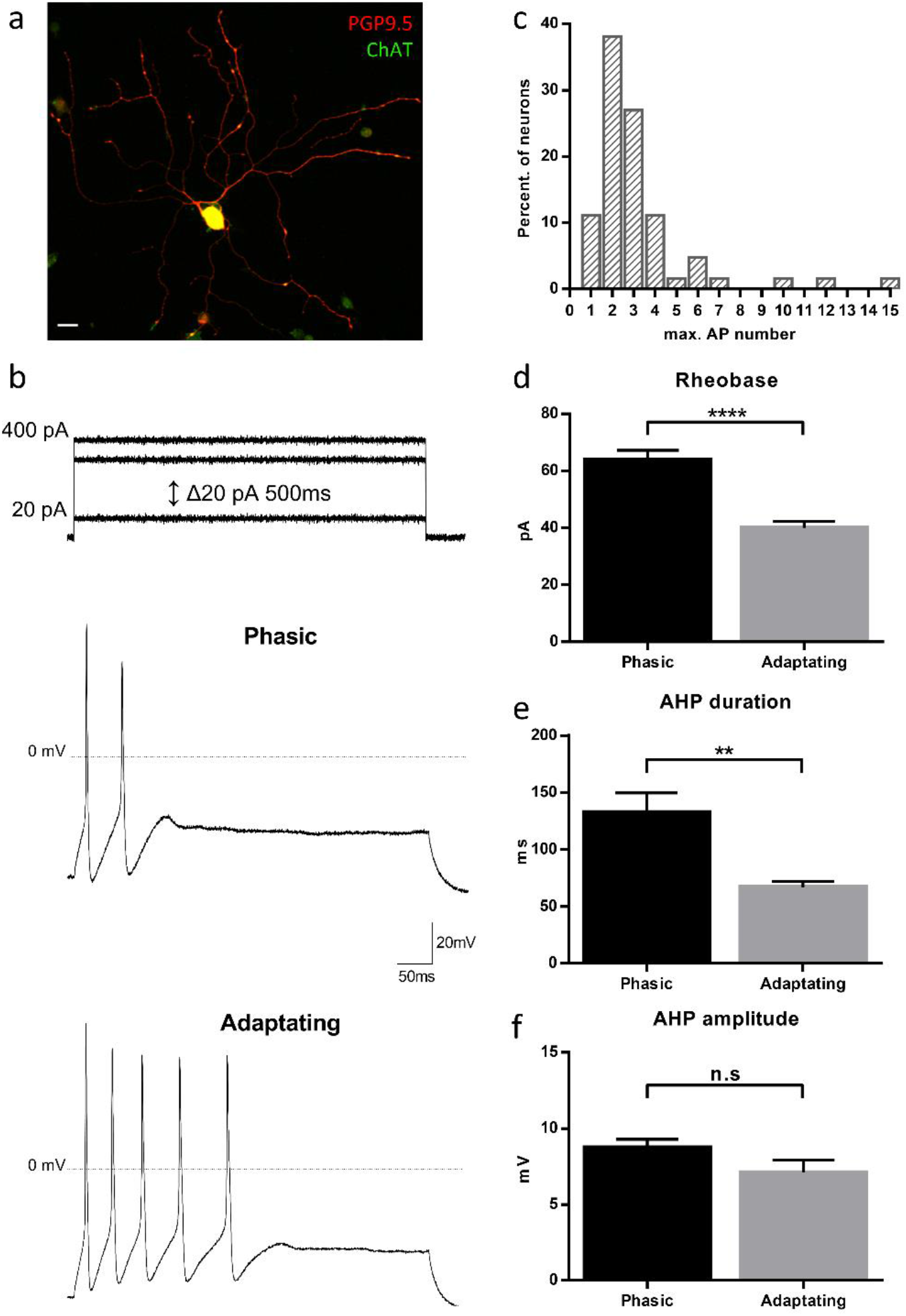
Action potential discharge profiles in mouse dissociated intracardiac neurons. **(a)** Mouse isolated cardiac neuron costained with the neuronal marker PGP9.5 and ChAT. **(b)** Maximum discharge of AP observed in two distinct neurons in response to 500ms depolarizing current injection from 20 to 400pA (Δ20pA). Upper trace represents AP obtained in a phasic neuron whereas lower trace was obtained in an adapting one **(c)** Distribution of the maximum number of AP recorded with the stimulation protocol presented in a. **(d)** Assessment of the rheobase of phasic (n=48) versus adapting neurons (n=14) (p<0,0001) **(e-f)** Assessment of the duration and amplitude of AHP in phasic (n=21) versus adapting (n=7) neurons with the stimulation protocol used in figure 5 (AHP duration p<0,01).

After 12 to 30h of culture, passive and active electrical membrane properties of isolated mouse cardiac neurons were assessed using the patch clamp technique in current clamp mode.

In our experimental conditions, no spontaneous firing activity was observed. Investigation of discharge characteristics revealed two different profiles. 77.4% of neurons showed limited firing activity (phasic neurons) characterized by a maximum of 1 to 3 action potentials (AP) at the onset of the stimulus, while 22.6% of neurons exhibited adapting firing behavior (fig.4b-c). Adapting neurons also showed a lower rheobase (40.0±2.3pA; n=14) than phasic neurons (64.0±3.3pA; n=48; p<0.0001, fig.4d). In addition, adapting neurons displayed a shorter afterhyperpolarization (AHP; 67.0±5.2ms (*adapting*) vs 132.8±17.0ms (*phasic*); p<0.01, fig.4e).

Upon brief injection of suprathreshold depolarizing current, intracardiac neurons were divided into two additional groups according to the presence or the absence of an AHP. 55% (30/55) of AP were followed by an AHP, while other neurons did not exhibit any (fig.5). Neurons with AHPs were characterized by a more depolarized resting membrane potential (RMP) (−57.4±0.9mV (*AHP*) vs -62.7±1.1mV (*no AHP*); p<0.001, fig.5b), a smaller AP amplitude (120.8±2.8mV (*AHP*) vs 137.5±2.1mV (*no AHP*); p<0.0001, fig.5c) and a smaller maximum rate of depolarization (147.9±7.8V/s (*AHP*) vs 189.2±5.6V/s (*no AHP*); p<0.001, fig.5d). It is noteworthy that these two newly defined types of neurons were not correlated to phasic or adapting patterns (p>0.05, Fisher’s exact test). Electrophysiological properties of mouse intracardiac neurons are summarized in supplementary table 1.

**Figure 5.**
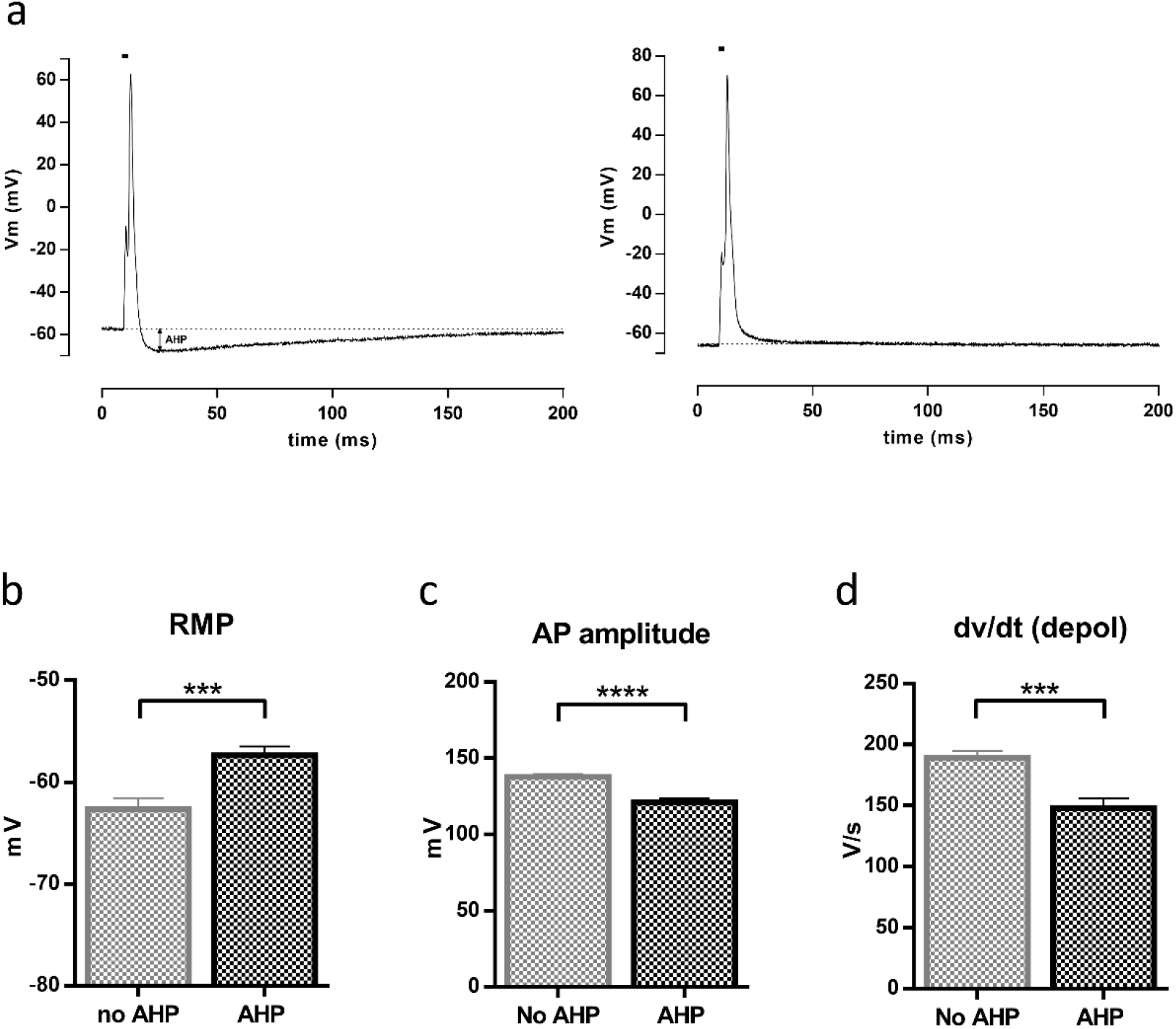
Action potentials properties in intracardiac neurons with or without AHP. AP with (left) or without (right) AHP recorded in response to a brief injection (2ms, black line) of suprathreshold depolarizing current **(a)**. Assessment of RMP **(b)** AP amplitude **(c)** and maximum rate of depolarization **(d)**. (***p<0,001, ****p<0,0001; no AHP (n=25); AHP (n=28)).

### Pharmacological response of mouse intracardiac neurons

Since the ICNS is under the dependence of several modulators^23^, we investigated the pharmacological response of isolated mouse cardiac neurons to known peripheral neuromodulators.

All tested neurons (n=28) exhibited membrane voltage and current responses to acetylcholine (Ach) perfusion, and the resulting inward current was sufficient to trigger AP (fig.6a). The superfusion of Ach resulted in the development of a large inward current, which amplitude decreased before the end of the drug application. This was probably due to channel desensitization. The peak current density evoked by Ach was similar in phasic vs adapting neurons, as well as in AHP vs no AHP neurons (fig.6b).

**Figure 6.**
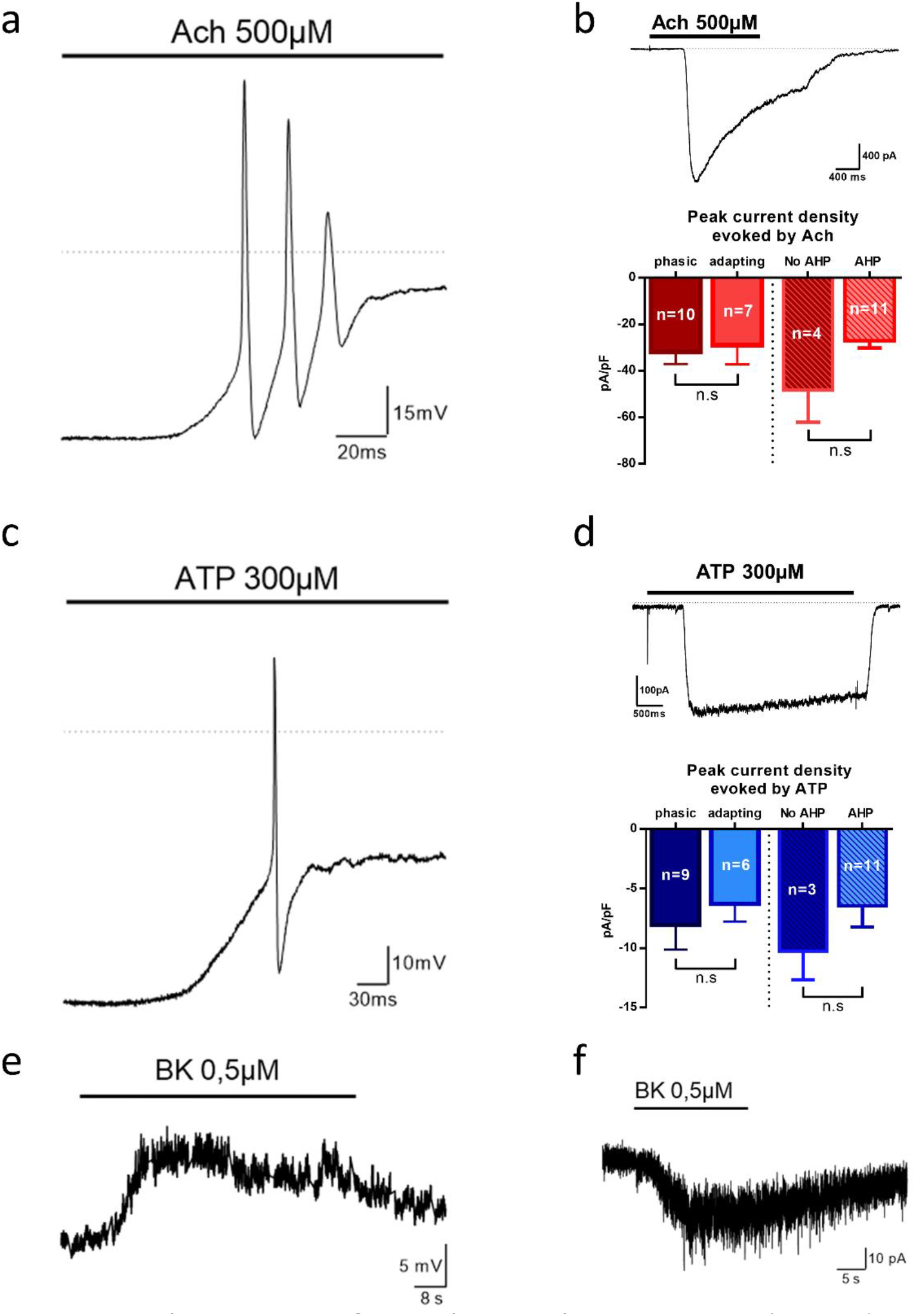
Pharmacological responses of mouse intracardiac neurons. Membrane voltage (left) and current (right) responses to Ach (500µM)**(a)**, ATP (300µM)**(b)** and BK (0,5µM)**(c)**.

The expression of several purinergic receptors has been described in rat cardiac neurons^24^ and adenosine triphosphate (ATP) and other purinergic compounds have been reported to modulate intracardiac neurons^23^. In our study, exogenous application of ATP was accompanied by a strong membrane depolarization in all tested neurons (n=17), resulting most of the time in AP firing (fig.6c). In voltage clamp, ATP induced a rapidly activating and sustained inward current, distinctive of purinergic receptor activation. There was no statistical difference between phasic and adapting neurons, as well as between AHP and no AHP neurons (fig.6d).

In our study, perfusion of bradykinin (BK) was able to slowly depolarize mouse cardiac neurons without eliciting AP (0.2µM: 8.4±1.1mV (n=7); 0.5µM: 13.0±2.4mV (n=8)). This was confirmed by the observation of a very slowly activating inward current of small amplitude after BK exposure in voltage clamp experiments (fig.6f).

## Discussion

This study presents for the first time a detailed report of anatomic, phenotypic, electrophysiological and pharmacological properties of the mouse ICNS.

### Cholinergic and catecholaminergic innervation of mouse heart

For the past thirty years, cardiac innervation and ICNS have been studied in a variety of species using tissue sections or whole-mount approaches. Although these techniques have brought a lot of information regarding the cardiac autonomic innervation, they essentially report superficial information and suffer from limitations because of sample damaging. The recent development of optical clearing techniques combined with advances in 3D imaging of large-scale specimen helped overcome these limitations, opening new opportunities to study global cardiac innervation.

The iDISCO clearing method allowed us to visualize and quantify in three dimensions the sympathetic and parasympathetic innervations of the mouse heart, thus enabling the determination of the exact location and number of cardiac ganglia in the whole non-sectioned murine heart. Our results are in accordance with Rysevaite et al.^25^, who reported a mean number of 19±3 ganglia. Since three dimensions analysis of ganglia was missing in the literature, we quantified ganglion size using a volumetric approach and found that cumulative ganglia volume is 0.02 mm^3^. We also developed an automatic pipeline in ImageJ to quantify the total length of nerve fibers innervating the heart. Our approach is complementary to the remarkable work done by Rajendran et al.^26^ who developed a clearing-imaging-analysis pipeline to assess diameter and orientation of nerve fibers innervating the mouse heart. This anatomical description of neuronal cardiac circuits will bring useful information to better understand the autonomic cardiac innervation and to identify abnormal cardiac innervation in pathological states, as already described in myocardial infarction or in cardiac autonomic neuropathy^27,28^.

### Neurochemical phenotype of mouse intracardiac neurons

So far, very little was known about the neurochemical diversity of intracardiac neurons in mouse, with only three different neuronal markers (ChAT, TH and nNOS) identified^2,7,8^. Our results confirm that almost all intracardiac neurons express the cholinergic marker ChAT, and that nearly 30% of them co-express ChAT and TH^7,8^. We also identified a small population of nitrergic neurons which is consistent with previous investigations that have reported such a phenotype in rat, guinea pig, rabbit and human^3–5^.

For the first time, we described the expression of four additional markers within somata of mouse intracardiac neurons, thus showing that this species is characterized by a relative phenotypic diversity, as reported in others mammals^3,5,10^. Indeed, we emphasized the existence of neurons expressing the calbindin, NPY and CART peptides and to a lesser extent glutamatergic neurons. A large part (46%) of mouse intracardiac neurons expressed the calcium binding protein calbindin. To date, only one study had identified such neurons in rat heart, where they represent only 7% of total neurons^3^. The expression of this protein has been reported in central and peripheral neurons, and is often used as a marker to discriminate different functional subpopulations of neurons, such as sensory neurons^29^. We also observed a lot of intra- and inter-ganglionic fibers, as well as terminals surrounding cell bodies that are immunoreactive for calbindin, suggesting that calbindin-expressing neurons may be crucial components of local circuits. Whether the expression of this protein is associated with any specialized function still remains to be determined.

Recently, glutamatergic neurons immunoreactive for VGLUT1, VGLUT2 and glutaminase, the synthetic enzyme for glutamate, have been found within the ICNS of rat^9^. In our study, we occasionally observed neurons immunoreactive for VGLUT2, yet without detecting any VGLUT1 immunoreactivity. Therefore we suggest that in mouse, glutamatergic intracardiac neurons may exist but do not represent a significant neuronal population.

The expression of a variety of neuropeptides also accounts for the neurochemical diversity of intracardiac neurons, especially in rodents^6,11^. However, apart from the description of sensory fibers immunoreactive for CGRP and SP, the expression of other neuropeptides had not yet been investigated in mouse^7^. Here, we highlighted the expression of two neuropeptides, NPY and CART, within mouse intracardiac neurons. The expression of NPY within these neurons is not surprising since it is known to be widely distributed in the autonomic nervous system and its cardiovascular effects have been well documented^30^.

CART peptide expression concerns only a small number of cardiac neurons in the guinea pig while it was observed in 46% of neurons in the rat^11,12^. This proportion is even greater in mouse, with 61% of neurons showing immunoreactivity for CART. This peptide has been extensively studied in the enteric nervous system, where it is expressed in many neurons, but experimental evidence elucidating its biological function is still lacking^31^. Further studies will be necessary to clarify its function within the cardiac context.

### Electrophysiological and pharmacological properties of mouse intracardiac neurons

The complex organization of the ICNS was further supported by the examination of its electrophysiological properties. Indeed, based on their electrical behavior, different subtypes of neurons have been identified within mammals, demonstrating that intracardiac neurons are forming a heterogeneous population^14,16,17^. Here, we report the first detailed investigation of passive and active electrical properties of mouse intracardiac neurons. By studying their firing activity we identified two distinct neuronal populations as observed previously^15,17,21^.While most neurons were classified as phasic due to limited firing activity, a small proportion of neurons were able to discharge more AP (adapting neurons). Phasic neurons have a significant higher rheobase as well as a higher AHP duration than adapting neurons, which confirms the existence of two types of neurons displaying different excitability features. Little is known about the different ionic channels expressed by intracardiac neurons, especially in the mouse. Thus, it would be interesting to closely investigate the molecular determinants of these two types of electrical behaviors.

It is important to emphasize that our results were not obtained from freshly isolated cardiac neurons. Hence, they may have been affected by cell culture conditions. However, it is worth mentioning that our results are consistent with those previously obtained *in situ*^32^.

A significant number of studies have revealed the diversity of neuromodulatory sources of intracardiac neurons which further suggests that they could play the role of integrative centers. In dog, neuronal activity is known to be regulated by mechanical and chemical stimuli^27^. Similarly, a variety of substances have been reported to modulate excitability of intracardiac neurons in rat and guinea pig^13,23,33^. However, the pharmacological modulation of mouse intracardiac neurons has never been investigated. Here, we showed that mouse cardiac neurons respond to Ach, ATP and BK, suggesting that they could be modulated by a variety of stimuli. In our experiments, BK induced a small inward current associated to a slight membrane depolarization. This indicates that BK receptors are present in mouse cardiac neurons but their stimulation was not sufficient in our conditions to trigger AP firing as observed in rat^33^.

## Conclusion

In conclusion, our study is the first detailed report providing phenotypic, electrophysiological and pharmacological characterization of mouse intracardiac neurons. Our results demonstrate that the mouse ICNS shares similar complexity to that of other species. ICNS complexity deserves consideration since there is growing evidence that ICNS plays an essential role in cardiac modulations and in the initiation and maintenance of cardiac arrhythmias. To this purpose, the use of transgenic mouse models will allow the functional investigation of neuronal phenotypes by expressing fluorescent reporter in specifically targeted neurons. Furthermore, the emergence of new genetic tools such as DREADDs and optogenetics opened new opportunities to control the activity of specific neurons within a global network. This represents a promising strategy to understand the role of targeted neurons in global cardiac modulation as described in the recent review of Scalco et al.^34^. However, these transgenic technologies are essentially available in mouse, where ICNS had been poorly characterized yet. Hence, our study is paving the way for future investigations using a combination of cre-mice systems and DREADDs/optogenetic tools in order to decipher the functional organization of the ICNS as well as its implication in pathological states such as arrhythmias.

## Author Contributions

G.L, J-F.F and A.C performed and analyzed the experiments. C.P conceptualized the imageJ pipeline. A.T and S.P performed technical imaging acquisition. G.L, C.P, J-F.F and A.C wrote the manuscript. A.C conceptualized and supervised the study.

## Acknowledgements

This work has benefited from the facilities of ImageUP platform (University of Poitiers) and the technical assistance of Anne Cantereau, Christophe Magaud and Cedric Bauer. We also thank Laura Batti for her help in the analysis of lightsheet images.

