## supplementary data for "Molecular and functional characterization of the mouse intracardiac nervous system"

**Supplementary table 1: electrophysiological properties of mouse intracardiac neurons.**

| **Parameters** | **Phasic** | **Adapting** | **No AHP** | **AHP** |
|---|---|---|---|---|
| Resting membrane potential (mV) | -60,0 ± 0,8 (n=48) | -59,8 ± 1,4 (n=15) | **-62,7 ± 1,1 (n=25)** | **-57,4 ± 0,9 (n=30)*** ** |
| Input resistance (MΩ) | 1392 ± 82 (n=30) | 1558 ± 103 (n=6) | 1365 ± 83 (n=20) | 1489 ± 122 (n=16) |
| Capacitance (pF) | 35,7 ± 1,6 (n=45) | 35,4 ± 3,0 (n=14) | 33,6 ± 1,9 (n=25) | 35,9 ± 2,1 (n=29) |
| Rheobase (pA) | **64,0 ± 3,3 (n=48)** | **40,0 ± 2,3 (n=14)****** | 64,4 ± 5,0 (n=25) | 54,7 ± 3,5 (n=30) |
| Spike half-width (ms) | 1,94 ± 0,05 (n=41) | 1,79 ± 0,1 (n=12) | 1,88 ± 0,07 (n=25) | 1,93 ± 0,07 (n=28) |
| AP amplitude (mV) | 128,1 ± 2,0 (n=41) | 130,5 ± 6,6 (n=12) | **137,5 ± 2,1 (n=25)** | **120,8 ± 2,8 (n=28)****** |
| dV/dt (depol.) (V/s) | 164,5 ± 6,0 (n=41) | 177,3 ± 14,7 (n=12) | **189,2 ± 5,6 (n=25)** | **147,9 ± 7,8 (n=28)*** ** |
| dV/dt (repol.) (V/s) | -84,1 ± 2,9 (n=41) | -97,7 ± 10,2 (n=12) | -90,8 ± 4,5 (n=25) | -83,9 ± 4,7 (n=28) |
| AHP duration (ms) | **132,8 ± 17,0 (n=21)** | **67,0 ± 5,2 (n=7)**** | | 8,4 ± 0,5 (n=29) |
| AHP amplitude (mV) | 8,5 ± 0,6 (n=25) | 7,1 ± 0,8 (n=8) | | 112,6 ± 14,4 (n=26) |

dV/dt: maximum rate of the depolarization or repolarization phase. AHP duration and amplitude were only calculated for neuron displaying AHP. **P<0,01; ***P<0,001. **** P<0,0001 Mann Whitney test.
