## supplementary material for "Molecular and functional characterization of the mouse intracardiac nervous system"

#### Animals

27 animals were used for this work :
- 3 mice for the iDISCO clearing approach
- 13 mice for immunohistochemistry
- 11 mice for the electrophysiological experiments

#### iDISCO heart clearing

Mouse hearts were stained and cleared using a modified iDISCO+ protocol[1,2]. Fixed hearts were dehydrated with a graded methanol series (50, 80 and 100% methanol in PBS, each for 1.5 h at room temperature (RT)) and bleached overnight with 6% H202 in methanol at 4 °C. After two washes in 100% methanol, hearts were gradually rehydrated (80, 50% methanol and PBS, 1h30 each).

Samples were blocked and permeabilized with PBS containing 0.2% gelatin 0.5% triton X-100 and 50mM sodium azide (PBSGT) for 4 days at RT. Primary antibodies (see supplementary table 1) were incubated during two weeks in PBSGT supplemented with 0.1% saponin at 37°C. Samples were then washed in PBSGT for 1 day followed by a 48 hours incubation with secondary antibodies. After another day of washing, samples were again dehydrated in methanol (20, 40, 60, 100 and 100% methanol, 1h each) and incubated overnight in a mixture of dichloromethane (DCM) – methanol (2:1) at RT. Samples were incubated in 100% DCM for 30mn and finally incubated and stored in DiBenzyl Ether.

Imaging was performed using a mesoSPIM system[3] at Wyss Center, Geneva. Briefly, the sample is illuminated by two digitally scanned light sheets coming from opposite directions. The excitation paths also contain galvo scanners for light-sheet generation and reduction of shadow artifacts due to absorption of the light-sheet. In addition, the beam waist is scanned using electrically tunable lenses (ETL, Optotune EL-16-40-5D-TC-L) synchronized with the rolling shutter of the sCMOS camera. This axially scanned light-sheet mode (ASLM) leads to a uniform axial resolution across the field-of-view (FOV). Emitted fluorescence is collected by high-numerical-aperture objectives (Olympus MVPLAPO 1 X – NA 0,25) and imaged on a digital camera (Hamamatsu ORCA-Flash 4.0).

Image processing and analysis was performed using Bitplane Imaris. Quantification of ganglionic volume was performed using the surface mode of the software based on ChAT staining. Ventricular innervation analysis was performed using ImageJ custom routines. Briefly, it consists in: First, discriminating the innervation from the background noise by thresholding the 3D image using a ‘maximum entropy’ algorithm[4]. The result is an image showing only the neural fibers network. Second, a ‘skeletonization’ that consists in reducing the diameter of all fibers to one voxel. This corrects the overestimation of the fiber volume in the image due to the point spread function, and the heterogeneity of illumination and optical density within the heart. The efficiency of these first parts of the analysis was validated by a systematic control of the overlap of neural network of a z-projection of a raw image and over-threshold fibers of a z-projection of a skeletonized image. Third, computing the length of the neural network using a custom version of the ImageJ plugin ‘analyse skeleton’[5].

**Immunohistochemistry**

After injection of heparin, mice were euthanized with an intraperitoneal injection of pentobarbital sodium (100 mg/kg). Hearts were quickly removed, washed in cold Tyrode solution and fixed in 4% paraformaldehyde for 24h at 4°C. After 3 washes in PBS, hearts were cryoprotected in 30% sucrose overnight at 4°C. Just before sectioning, samples were embedded in OCT tissue compound (Tissue Tek) and frozen in cold ethanol. Sections 40–50 µm thick were cut with a cryostat and were collected on microscope slides (Adhesion slides, Menzel Gläser, SuperFrost® Plus, Thermo scientific). Sections were washed 3 times with PBS and permeabilized with 0.5% TritonX-100/1% BSA in PBS for 2h at RT. Incubation with primary antibodies (supplementary table A) was performed overnight at 4°C. After 3 washes with PBS, sections were incubated with secondary antibodies for 3h in the dark at RT. Nuclei were stained with DAPI and sections were mounted in mowiol mounting medium. Images were acquired using a confocal laser scanning microscope (FV3000 Olympus). As a control, images obtained after incubation with the secondary antibodies in the absence of primary antibodies did not elicit any labelling. Quantification was performed by manual cell counting using NIH ImageJ (Bethesda, Maryland, USA). For each marker, quantification was done with sections coming from at least two distinct animals. All micrographs are a projected confocal Z-series from sectioned material.

**Neuron dissociation**

Mice were injected with heparin and euthanized with pentobarbital sodium (100 mg/kg). Hearts were quickly removed and washed in cold HBSS solution. Fat pads located between both atria were dissected, cut into small pieces and dissociated in 2 mL HBSS containing 3 mg/mL collagenase type II (Worthington), 7.5mg/mL dispase II and 0.25 mg/mL DNase I for 30 mn at 37°C. This was followed by an additional incubation in 2 mL trypsin-EDTA 0.25% supplemented with 0.25 mg/mL DNase I for 35 mn at 37°C. After two washes in culture media, cells were gently triturated with fire-polished Pasteur pipettes coated with SVF and plated on laminin-coated 35 mm Petri dishes. Cells were maintained in Neurobasal-A medium supplemented with 2mM L-glutamine, B27 supplement, 5% horse serum and 1% penicillin/streptomycin in a humidified chamber at 37°C with 5% $CO_2$.

**Electrophysiology**

Passive and active electrical membrane properties of isolated cultured neurons were determined using the whole-cell configuration of the patch clamp technique in current clamp mode. For pharmacological studies, drugs were perfused using a gravity perfusion system (Ala Scientific Instruments) and ionic currents were assessed using the voltage clamp mode at a holding potential of -60mV. Recordings were carried out at RT within 30 hours following neurons isolation. Patch electrodes (≈4 MΩ) were pulled from glass capillaries (PG150T-7.5, Harvard Apparatus, Les Ulis, France) using a vertical micropipette puller (Narishige, Tokyo, Japan). The patch pipettes were filled with (mM): 130 K-gluconate, 10 KCl, 1 MgCl2, 10 HEPES, 1 CaCl2, 5 EGTA, 2 Mg-ATP, 10 Na2-phosphocreatine and 0.3 Na-GTP, (pH adjusted to 7.2 using KOH). The bath solution contained (mM): 150 NaCl, 5 KCl, 1 MgCl2, 1.5 CaCl2, 10 glucose, and 10 HEPES, (pH adjusted to 7.4 using NaOH). Liquid junction potential was +16 mV and was not corrected. Recordings were made with an Axopatch 200B amplifier (Molecular devices, San Jose, California, USA) with a 5-kHz low-pass filter. Data were sampled at 20 kHz and digitized by a Digidata 1550 B (Molecular devices, San Jose, California, USA). Data acquisition and analysis were performed using pClamp software (v11, Molecular devices, San Jose, California, USA).

Input resistance was determined by measuring voltage changes evoked by injection of hyperpolarizing current (from -10 pA to -40 pA; in -10 pA increments). Discharge characteristics were determined by injecting depolarizing current of increasing amplitude (from 20 to 400 pA ; 20 pA increment) during 500 ms. AP properties were determined upon brief injection (2 ms) of suprathreshold current. AP amplitude was measured as the difference between the AP peak amplitude and the resting membrane potential. Spike half-width was calculated as the AP duration measured at 50% of its amplitude.

### Chemicals

PACAP27, VIP and Bradykinin was obtained from Bachem (Bubendorf, Switzerland), oxotremorine-M from Tocris Bioscience (Bristol, United Kingdom). Neurobasal A medium and B27 supplement was supplied by ThermoFisher Scientific (Villebon sur Yvette, France). Unless stated, all others chemicals was obtained from Sigma-Aldrich (Lyon, France).

**Supplementary Table A :** Primary and secondary antisera used within this study.

| Primary Antibody | Host Species | Dilution | Catalogue Number | Supplier |
|---|---|---|---|---|
| Calbindin D28k | Rabbit | 1:1200 | CB38 | Swant |
| CART (55-102) | Rabbit | 1 :1000 | H-003-62 | Phoenix Pharmaceuticals |
| CGRP | Goat | 1:300 | Ab36001 | Abcam |
| ChAT | Goat | 1:150 | AB144P | Merck Millipore |
| MAP2 | Rabbit | 1:600 | AB5622 | Merck Millipore |
| nNOS | Rabbit | 1:500 | AB5380 | Merck Millipore |
| NPY | Rabbit | 1:500 | Ab10980 | Abcam |
| PGP9.5 | Guinea pig | 1:200 | Ab10410 | Abcam |
| PGP9.5 | Rabbit | 1:2000 | AB1761-I | Merck Millipore |
| Somatostatin | Rat | 1:200 | sc-47706 | Santa cruz |
| Substance P | Guinea pig | 1:500 | Ab10353 | Abcam |
| TH | Chicken | 1:500 | Ab76442 | Abcam |
| TH | Rabbit | 1:400 | AB152 | Merck Millipore |
| VIP | Rabbit | 1 :500 | 20077 | Immunostar |
| VGLUT2 | Guinea pig | 1 :400 | 135404 | Synaptic Systems |
| **Secondary Antibody** | | | | |
| Anti-chicken (FluoProbe594) | Donkey | 1:300 | FP-SD1110 | Interchim |
| Anti-goat (AF488) | Donkey | 1:300 | A-11055 | ThermoFisher Scientific |
| Anti-goat (AF555) | Donkey | 1:300 | A-21432 | ThermoFisher Scientific |
| Anti-goat (AF647) | Donkey | 1:300 | A-21447 | ThermoFisher Scientific |
| Anti-guinea pig (AF488) | Goat | 1:300 | A-11073 | ThermoFisher Scientific |
| Anti-guinea pig (AF647) | Donkey | 1:200 | 706-605-148 | Jackson Immunoresearch |
| Anti-rabbit (AF488) | Chicken | 1:300 | A-21441 | ThermoFisher Scientific |
| Anti-rabbit (AF555) | Donkey | 1:300 | A-31572 | ThermoFisher Scientific |
| Anti-rabbit (AF647) | Donkey | 1:300 | A-31573 | ThermoFisher Scientific |
| Anti-rat (AF488) | Goat | 1:300 | A-11006 | ThermoFisher Scientific |
